# The 16S rRNA Gene Is Not a Precise Classifier for *Luteibacter* Species

**DOI:** 10.1101/2022.07.18.500475

**Authors:** David A. Baltrus, Morgan E. Carter

## Abstract

*Luteibacter* species are found throughout agricultural and plant associated microbial communities, and have largely been identified and classified through comparisons of the 16S rRNA genes. Through comparisons of 16S classifications with whole genome phylogenies and ANI, we highlight a somewhat unique situation whereby *Luteibacter pinisoli* and *Luteibacter* sp. 9143 would be classified as the same species using 16S rRNA sequences but are clearly differentiated by these other metrics. We present this case as an outlier, but also as an example for the challenges of classification solely using 16S rRNA gene sequences.

## Introduction

The 16S rRNA gene was chosen and is still often used by a majority of research groups as a “barcode” marker to classify species within bacterial communities because it possesses a variety of qualities that are useful in this context (Johnson et al., 2019; Church et al., 2020). This choice has profoundly influenced our modern understanding of microbial community structure and dynamics, and is intricately interwoven into the foundations of microbiome research. As such, the 16S rRNA locus has become a go-to marker for rapid classification of bacterial strains (Srinivasan et al., 2015; Church et al., 2020). However, this situation creates assumptions that are problematic in a minority of cases, like the one described below, when the 16S rRNA locus does not reflect “true” genome level similarity but is still used for identification.

The 16S rRNA locus possesses numerous characteristics that make it an exquisite barcode for characterizing the microbial communities (Johnson et al., 2019). It is a critical component of the ribosomal translation complex and thus is present in all known bacterial isolates and genomes. It is relatively long (∼1500bp) and thus contains multiple regions that can accumulate genetic diversity, which ultimately enables discrimination across taxa. However, it also contains conserved sites bracketing these regions of diversity, which enable the construction of degenerate primer pairs that can amplify these regions in a targeted way (reference). There also remains an overarching idea that, given its conserved and essential status within an important protein-RNA complex, it has little potential for recombination of the 16S locus across strains. For the most part, studies have largely borne this idea out: the 16S locus appears to be exchanged across strains at very low rates relative to other genes, but recombination and horizontal gene transfer has been demonstrated in (to this point) rare cases (Kitahara & Miyazaki, 2013; Tian et al., 2015).

Based on initial successes classifying strains using the 16S rRNA gene, numerous analytical pipelines have been developed to categorize microbial communities using this locus and multiple databases (e.g. SILVA (Quast et al., 2013)) have been created to provide an overarching and somewhat reliable structure for classification by 16S rRNA gene sequences. These discussions dovetail with taxonomic arguments and assessments about “appropriate” levels of divergence in 16S rRNA sequences, with current convention being that sequences sharing >97% identity (<3% divergence) classified as different operational taxonomic units (OTUs) across papers and taxonomically as different species if comparing cultured isolates (Nguyen et al., 2016; Edgar, 2018). The 97% threshold was originally chosen to represent a level of sequence similarity at the 16S rRNA gene that reflected 70% thresholds for whole genome DNA-DNA hybridization between genomes containing these genes (Stackebrandt & Goebel, 1994).

However, a variety of weaknesses have also arisen in the context of using the 16S rRNA gene for characterization of cultured and uncultured microbial communities (Rajendhran & Gunasekaran, 2011). First, there is no universal primer set that works in an unbiased way across all bacterial types and lineages; thus, every study has the potential for either missing the identification of lineages completely or miscalculating the frequencies of bacterial types (Hong et al., 2009; Pinto & Raskin, 2012; Brooks et al., 2015). This is as true of the 16S rRNA locus as it would be for any other gene found within the genome. Furthermore, even though an abundance of studies have suggested that the horizontal transfer of the 16S rRNA gene occurs relatively rarely if at all, some studies have highlighted the potential for exchange of parts of the ribosome (Tian et al., 2015; Miyazaki & Tomariguchi, 2019). Lastly, even though the 16S rRNA gene is a useful tool for classification of some species and lineages, there are other situations that have demonstrated genotypic and phenotypic variability across bacterial species that possess identical 16S rRNA gene sequences (Jaspers & Overmann, 2004). This is not a problem of mischaracterization by the 16S rRNA gene, however; it is a problem of lack of sufficient genetic diversity within the 16S rRNA gene sequence to accurately discriminate between relevant phenotypic units of bacteria. Conversely, there also exist documented instances where intragenomic divergence across multiple 16S rRNA gene copies within a single genome leads to incorrect classification and measurements of community level diversity (Louca, Doebeli & Parfrey, 2018).

Here we present a situation where the 16S rRNA gene sequence does not accurately classify relevant taxonomic groupings in that sequences of the 16S rRNA gene clearly mischaracterize relationships between strains within the genus *Luteibacter* compared to data from other loci as well as whole genome information. It is likely that this situation is due to a combination of relatively low levels of divergence in the 16S rRNA gene across *Luteibacter* compared to other parts of the genome as well as convergence in sequence in the 16S locus between distinct species. While we largely believe that this case is an outlier compared to the overwhelming data from a variety of bacterial taxa, we view this case as a way to highlight some of the caveats associated with relying on single locus data when inferring phylogenetic relationships.

## Materials and Methods

### Gene and Genome sequences

Individual gene sequences used in this manuscript were collected from the Joint Genome Institute’s Integrated Microbial Genome and Microbiome Portal (JGI IMG) but are also available through NCBI GenBank. Relevant accession numbers and strain information listed in Table 1. Outgroup strains were chosen to represent *Dyella* and *Xanthomonas* species, as these are consistently shown to be closely related to *Luteibacter* species using a variety of methods. For *Luteibacter* strains 9133, 9135, 9143, 9145, we have also deposited full length 16S rRNA gene sequences at NCBI, respectively, at accession numbers: ON786549.1, ON786546.1, ON786547.1, and ON786548.1.

**Table 1.**
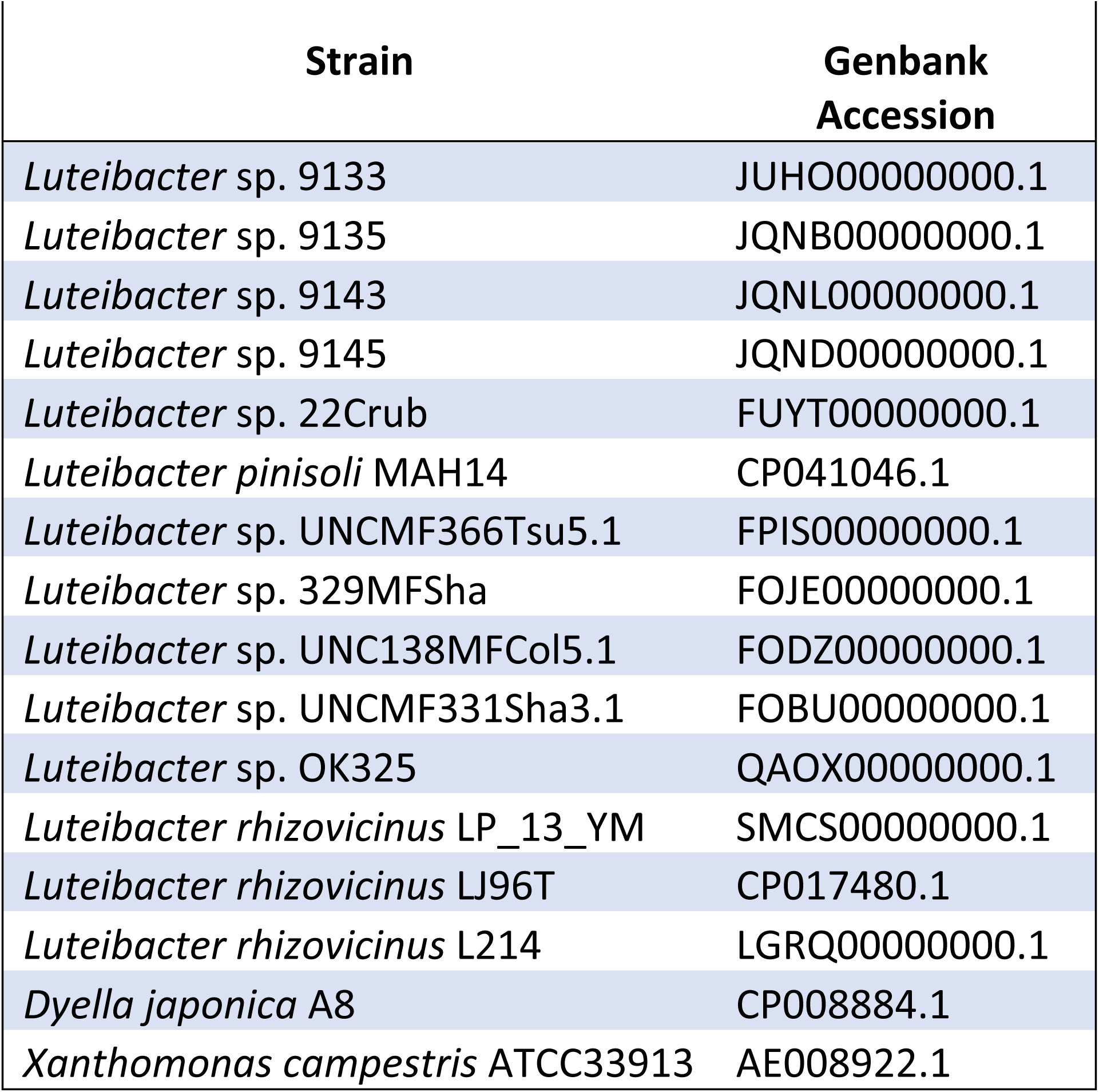
Genome Accessions of Strains Used in this Study

### Phylogenetic Inference and ANI Calculation

Gene sequences for full length 16S rRNA loci were obtained from JGI IMG for all strains in Figure 1, and were aligned and visualized for this figure using Geneious prime v2020.2.4. Aligned sequences can be found in the supplemental file at Figshare at <u>10.6084/m9.figshare.20333316</u>. For phylogenetic inference based on 16S rRNA genes, sequences were aligned using ClustalX with default parameters (Thompson, Gibson & Higgins, 2002). Ends of the sequence alignments were trimmed so that alignments from all sequences started and stopped at the same position. Bayesian phylogenies were inferred for 16S rRNA gene sequences using MrBayes 3.1 without priors (Ronquist & Huelsenbeck, 2003). Within MrBayes, sequences were run for 1,000,000 generations with 250,000 generations as a burn in and using GTR and invgamma for rates. Rates were chosen as the best fit by ModelTest-ng (Darriba et al., 2020).

**Figure 1:**
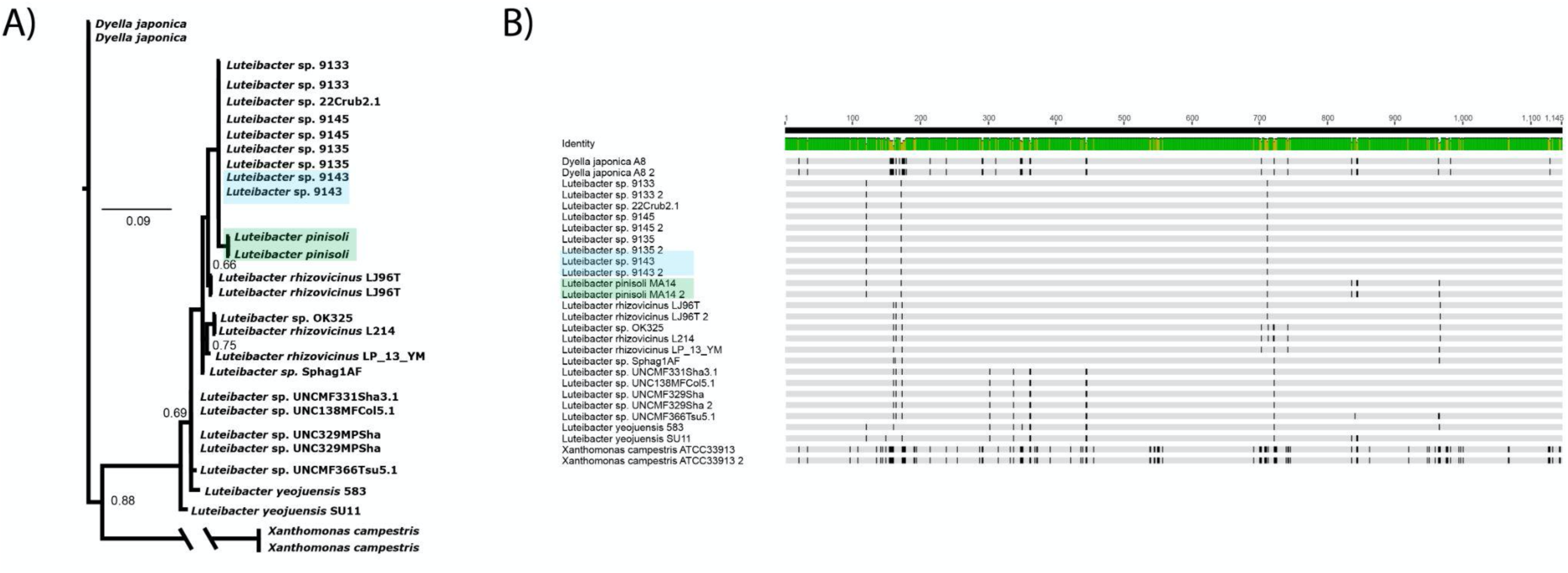
Similarity in 16S rRNA Gene Sequences Between *Luteibacter pinisoli* and *Luteibacter* sp. 9143. A) A Bayesian phylogeny inferred using nearly full-length 16S rRNA gene sequences from *Luteibacter* strains and related genera and species. Posterior probabilities of nodes with <95% support are shown. B) Alignments of nearly full-length rRNA gene sequences from *Luteibacter* strains and related species, built using ClustalW and visualized using Geneious. The top line reflects identity at each position, with green displaying identical sequences and yellow displaying divergent sites. Dark lines in each alignment indicate differences in nucleotide sequence from the consensus. *Luteibacter* sp. 9143 is highlighted in blue in both figures and *Luteibacter pinisoli* is highlighted in green in both figures.

The bacterial phylogenomic tree was produced with GToTree v1.6.31(Lee, 2019), using the prepackaged single-copy gene-set for bacteria. Briefly, prodigal v2.6.3 (Hyatt et al., 2010)was used to predict genes on input genomes provided as fasta files. Target genes were identified with HMMER3 v3.2.2 (Eddy, 2011), individually aligned with muscle v5.1 (Edgar, 2021), trimmed with trimal v1.4.rev15 (Capella-Gutiérrez, Silla-Martínez & Gabaldón, 2009), and concatenated prior to phylogenetic estimation with FastTree2 v2.1.11 (Price, Dehal & Arkin, 2010). TaxonKit (Shen & Ren, 2021)was used to connect full lineages to taxonomic IDs.

Whole genome phylogenetic inference was also carried out using the RealPhy server (Bertels et al., 2014), where *Xanthomonas campestris, Luteibacter* sp. 9143, and *Luteibacter pinisoli* were used as references. ANI calculator (Rodriguez-R & Konstantinidis, 2014) using default parameters was used to generate ANI data for each strain pair shown in the matrix in Figure 3, visualized with R v4.1.2 (R Core Team, 2016).

## Results and Discussion

When *Luteibacter pinisoli* was originally described and named as a species, sequence fragments from the type strain isolated from the roots of *Pinus koraiensis* in Korea placed this isolate as a new species within the genus *Luteibacter*, but outside of previously circumscribed species such as *L. rhizovicinus* (Akter & Huq, 2018). Phenotypic traits were concordant with this assessment. However, sequence fragments from *L. pinisoli* are nearly identical to a suite of unnamed strains (sp. 9143, sp. 9145, sp. 9135, sp. 9133) isolated as endohyphal bacteria found in association with a fungus in the United States (Durham, NC), referred to as *Pestalotiopsis* sp. 9143 (Hoffman & Arnold, 2010). We have since assembled a complete genome sequence for *L. pinisoli* (Baltrus et al., 2019) and for the suite of endohyphal strains possessing nearly identical sequences over their 16S rRNA gene fragments (Baltrus et al., 2017). There is high similarity in sequence over the entire length of the 16S rRNA gene locus for *L. pinisoli* and strain 9143 (representative of the currently unnamed clade), with decreasing levels of similarity found between this pair and other named species like *L. rhizovicinus* (Figures 1A, 1B). Overall, across the genus *Luteibacter*, there is very little divergence in the 16S rRNA gene (with full length sequence comparisons displaying >98% identity), (Figure 1A, 1B and Supplemental File 1). Moreover, phylogenies inferred from entire gene sequences of the 16S rRNA gene for *Luteibacter* species place *L. pinisoli* and 9143 as sister clades, although confidence intervals across the tree are relatively low due to the lack of informative sites (Figure 1A). Therefore, by 16S rRNA sequence alone, and using currently recognized conventions, the currently unnamed strains (9143, 9145, 9133, 9135) should be designated as members of *L. pinisoli*. It is important to note that relationships between these strains inferred from the 16S rRNA gene alone are consistent regardless of which genomic copy is used, as sequences found within the same genome are identical.

In contrast to the results from 16S rRNA gene classification, taxonomic assessment of *Luteibacter* strains using a variety of independent metrics strongly suggests that *L. pinisoli* clearly forms a separate taxonomic clade from the currently unnamed strains typified by strain 9143. (Figs. 2A, 2B, and 3). Phylogenies inferred using numerous conserved proteins as well as single nucleotide variants clearly differ from those inferred using the 16S rRNA gene results (Figures 2A and 2B). Instead of *L. pinisoli* forming a monophyletic clade including *Luteibacter* sp. 9143, whole genome metrics supported with high confidence suggest that *L. pinisoli* is clearly divergent from *Luteibacter* sp. 9143 and instead forms its own clade to the exclusion of *Luteibacter* sp. 9143 and related strains as well as additional *Luteibacter* strains and species. Moreover, ANI comparisons clearly show *L. pinisoli* to be quite divergent from *Luteibacter* sp. 9143 and that the ANI between *L. pinosoli* and multiple strains from this clade are <85% (Figure 3). Thus, phylogenetic relationships inferred using only the 16S rRNA gene are in conflict with those inferred using multiple different types of classifiers including whole genome information. This situation is distinct from others involving lack of resolution of the 16S rRNA gene to differentiate between extensive phenotypic diversity across similar strains, because whole genome phylogenies actually suggest that *L. pinisoli* is distantly related to *Luteibacter* sp. 9143. We highlight that this is not simply the case of too little resolution at the 16S locus compared to other phylogenetically relevant sites as numerous strains can be classified as intermediate between *L. pinisoli* and *Luteibacter* sp. 9143 according to conserved protein, whole genome metrics, and broader classifiers such as ANI. Specifically, we highlight that *Luteibacter* sp. 9143 and *Luteibacter pinisoli* share >98% sequence identity throughout the 16S rRNA locus but an ANI comparison of <83%. This ANI value is far below the convention of <95% ANI for classification as different bacterial species. That genomes for these strains show relatively more divergent 16S rRNA gene sequences than those found in *L. pinisoli/Luteibacter* sp. 9143 is definitive data supporting the idea of improper classification by 16S rRNA gene sequences. While we recognize that there are also a variety of challenges associated with whole genome classification (Young & Gillung, 2020), we highlight that whole genome level analyses presented herein are performed using multiple different tools and datasets and that phylogenetic inference is independent from ANI calculations.

**Figure 2:**
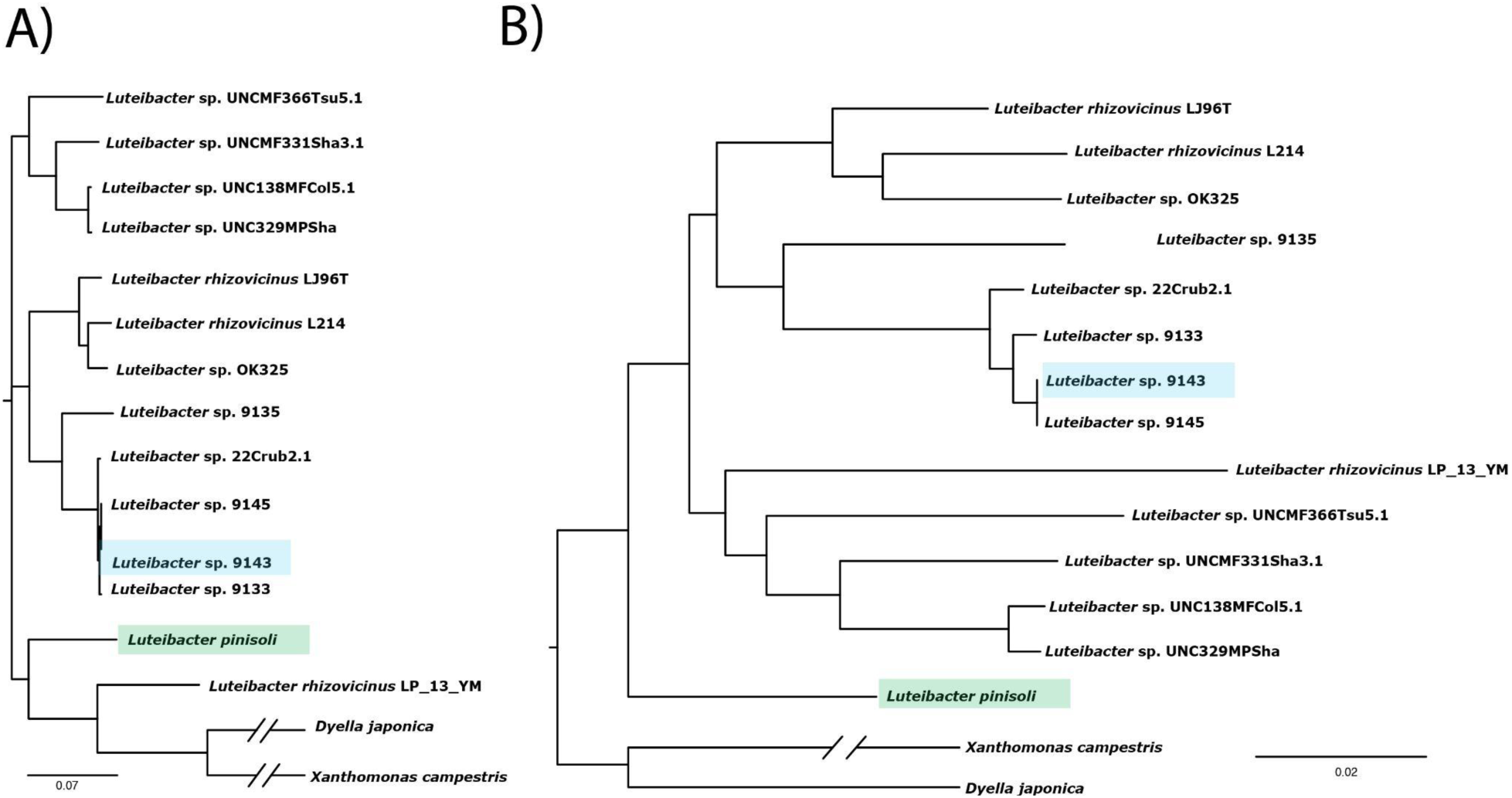
Phylogenies of *Luteibacter* and Related Genera inferred Using Whole Genome Information. A) A maximum likelihood phylogeny inferred using single copy genes conserved across genomes, using the program GToTree B) A maximum likelihood phylogeny inferred using informative genomic variants and built using RealPhy. *Luteibacter* sp. 9143 is highlighted in blue in both figures and *Luteibacter pinisoli* is highlighted in green in both figures.

**Figure 3:**
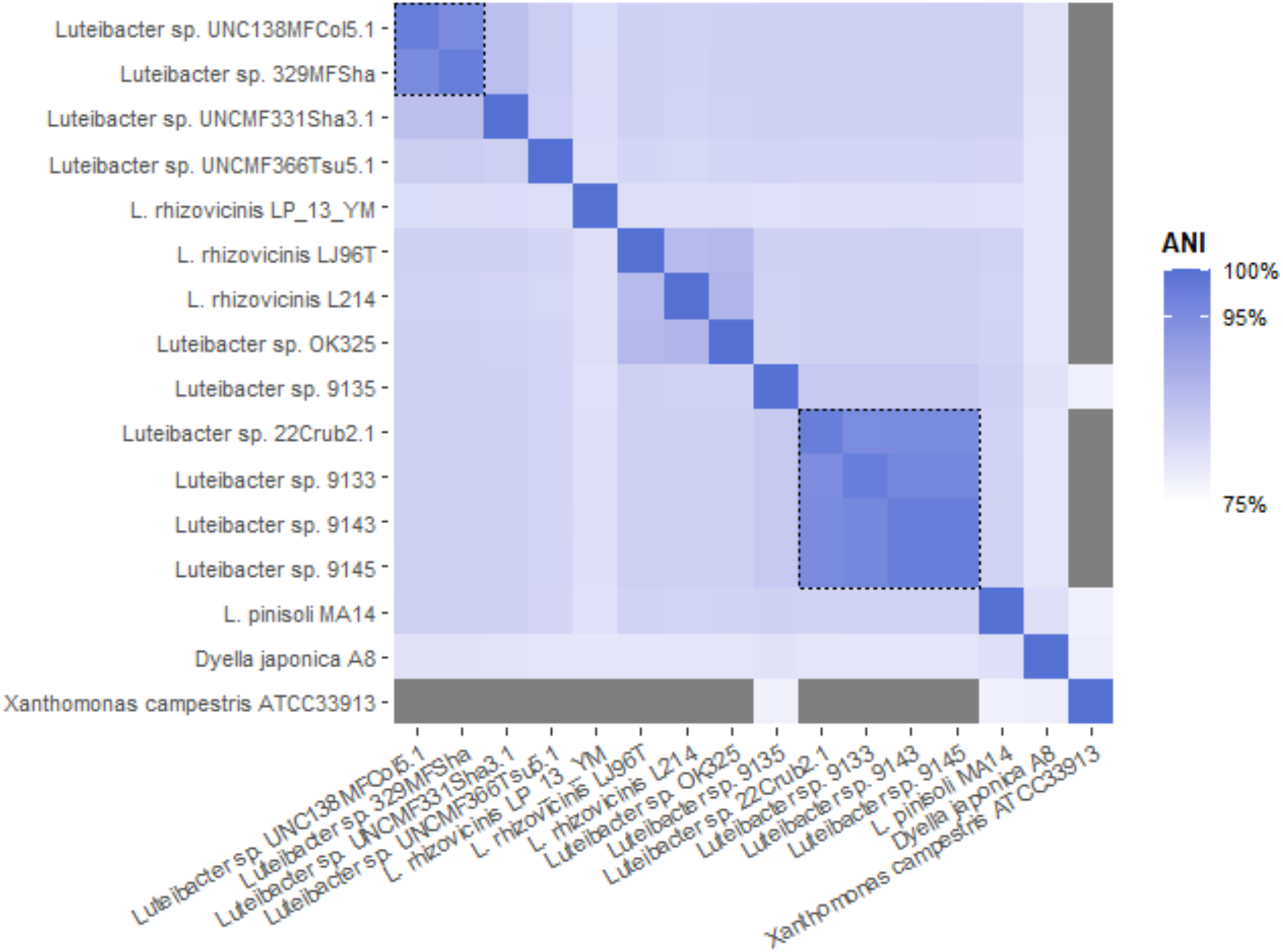
ANI Comparisons Across *Luteibacter* and Related Strains and Species. ANI calculator was used to compare whole genome sequences of strains shown below. ANI similarity is color coded as per the legend, with blue representing 100% and lighter shades of blue representing decreasing similarity in ANI. Grey boxes indicate ANI values <75%. Dotted lines indicate groupings where ANI values >95%, the typical threshold for species distinctions.

The main reason that the 16S rRNA gene is not precise for classification within *Luteibacter* is simply that there is very little divergence in this locus across the genus, with similarity in sequence for this locus between any given pair of *Luteibacter* isolates being >98% (Figures 1A, 1B). In addition, as one can see in the alignments in Figure 1B, there exists additional levels of convergence of sequence for this locus between the strain labeled as *pinisoli* and that labeled as sp. 9143. Two possible explanations are possible for this convergence. First, a section of the 16S rRNA locus could have been horizontally transferred between these two clades of *Luteibacter*. When the 16S rRNA gene is split into “variable” sections, it is noteworthy that V3-V6 appear to support close relationships between *L. pinisoli* and strain 9143, but V8 does not (Supplemental Figure 1). However, there are currently too few strains isolated and sequenced from within and closely diverged from the *Luteibacter* sp. 9143 clade to clearly identify areas of recombination given the relatively sparse number of polymorphisms within this locus. It is also possible that there have been a series of convergent nucleotide substitutions within the 16S rRNA gene in the genome of *L. pinisoli* to match those within *Luteibacter* sp. 9143. There currently exists no mechanism to explain such a trend, aside from linkage of mutations at multiple sites within rRNA molecules due to selective pressures on secondary structure, but we cannot rule it out with available data at present. Indeed, whatever the mechanism for convergence in 16S rRNA sequence between *L. pinosoli* and *Luteibacter* sp. 9143, it has to also explain how both copies of the 16S rRNA loci within both genomes display the same trends.

Overall, we have demonstrated that the 16S rRNA gene is not a faithful classifier for *Luteibacter* species. Although this locus does work well to classify some clades and strains, we have shown that strains *L. pinisoli* and *Luteibacter* sp. 9143 clearly form separate clades when assessed by multiple whole genome metrics despite nearly identical 16S rRNA gene sequences. We note that this situation does not solely appear to be simply explained by a lack of resolution in the 16S rRNA locus itself, although there are relatively few polymorphisms compared to other loci, but that there are convergent nucleotide changes that artificially cause clustering between *L. pinisoli* and strains like 9143. At present we cannot differentiate between scenarios involving convergent substitution of nucleotides or horizontal gene transfer of the 16S rRNA gene itself. Taken together, whole genome phylogenies as well as divergence in the ANI data suggest that *Luteibacter* sp. 9143 and *L. pinisoli* are members of separate species, even though they have nearly identical 16S rRNA genes.

